# Evidence for the Transient Presence of Atypical Astrocytes in Mice Following a Single, Closed-Head Mild Traumatic Brain Injury

**DOI:** 10.1101/2025.09.16.676600

**Authors:** Jesse B. Blackman, Rebecca Krauss, Sadi Quiñones, Panorea Tirja, Mary E. Sommer, Carmen Muñoz-Ballester, Farzad Noubary, Moritz Armbruster, Stefanie Robel, Trent Anderson, Chris G. Dulla

**Affiliations:** Department of Neuroscience, Tufts University School of Medicine, 136 Harrison Avenue, Boston, Massachusetts, 02111, USA; Neuroscience Program, Tufts Graduate School of Biomedical Sciences, Tufts University, 136 Harrison Avenue, Boston, Massachusetts, 02111, USA; Department of Cell, Developmental and Integrative Biology, University of Alabama at Birmingham, 1900 University Boulevard, Birmingham, Alabama, 35294, USA; Department of Biological Sciences, University of Maryland Baltimore County, 1000 Hilltop Circle, Baltimore, Maryland, 21250, USA; Department of Public Health and Health Sciences, Bouvé College of Health Sciences, Northeastern University, 360 Huntington Avenue, Boston, Massachusetts, 02115, USA; Department of Basic Medical Sciences, College of Medicine, University of Arizona, 475 North 5th Street, Phoenix, Arizona, 85004, USA

**Keywords:** Atypical astrocytes, astrogliosis, glutamate transporters, mild TBI, brain injury, mouse model

## Abstract

Mild traumatic brain injury (mTBI) affects roughly 42 million people each year, causes a variety of physical, behavioral, and cognitive symptoms, and increases the risk for developing neurological disorders, including post-traumatic headache (PTH) and Alzheimer’s disease (AD). Multiple molecular and cellular changes occur following mTBI; here we focus on astrocytes - cells that respond to brain injury and are critical to maintaining neuronal and circuit homeostasis. While some astrocytes become reactive after mTBI, others adopt an ‘atypical’ state characterized by the loss of multiple functional astrocyte proteins, including glutamate transporters (GLT-1, GLAST) and ion channels (Kir4.1), without upregulation of prototypical reactive astrocyte markers (glial fibrillary acidic protein [GFAP]). Previous studies have shown that repeated mTBI causes atypical astrocytes (AtAs) that can persist for months, but we know much less about whether a single mTBI causes similar astrocyte phenotypes. To address this, we employed a closed-head mild traumatic brain injury (chmTBI) model in male and female mice and quantified the abundance of AtAs both acutely (3-days) and chronically (1-month) after a single injury. We found that 3-days after chmTBI, AtAs were present in areas subject to blunt force trauma (BFT), consistent with previous reports, as well as in other brain regions presumably affected by diffuse injury. One month after chmTBI, however, the proportion of AtAs was similar between chmTBI and sham injured mice, thereby suggesting AtAs do not persist long term in this model. Consistent with previous studies, this chmTBI model did not induce significant GFAP-positive reactive astrocytes as assayed using immunohistochemistry, at either timepoint. Overall, we show an increase in AtAs 3-days after a single chmTBI that returns to sham levels when examined 1-month after injury. This suggests that after a single mTBI, AtAs are present but do not persist long term, unlike in repeated mTBI where AtAs persist for months after injury.

## Introduction

Traumatic brain injury (TBI) is a disruption to brain function caused by an external physical trauma, is a leading cause of disability and mortality, affects more than 42 million people globally^1^, and is classified based on injury severity. Mild traumatic brain injury (mTBI) is the most common form, accounting for approximately 80% of all TBI cases^2^. mTBI contributes to a variety of neurological and psychiatric dysfunctions (e.g. headaches, attentional problems, and anxiety^3–6^) and increases patient risk of developing disorders such as post-traumatic headache, major depressive disorder, and Alzheimer’s disease (AD)^6–8^. Following mTBI, multiple cellular and molecular changes unfold, some of which are protective, while others contribute to secondary injury^9, 10^ and include blood brain barrier (BBB) disruption, neuroinflammation, and excitotoxicity^9, 11, 12^ - processes in which astrocytes play a critical role. Astrocytes are an abundant type of glia in the central nervous system that help maintain brain function by regulating neurotransmitters and extracellular ions, providing metabolic support of neurons, contributing to BBB structure and function, and modulating inflammation^13^. Following injury, astrocytes can become reactive, changing their gene expression, structure, and function^14^.

Recent studies have shown that astrocytes can adopt an ‘atypical’ phenotype following single or repeated mTBI (rmTBI) induced by a modified Marmarou weight-drop model^15, 16^. Atypical astrocytes (AtAs) are identified by a lack of multiple astrocytic functional proteins, including glutamate transporter 1 (GLT-1), glutamate/aspartate transporter (GLAST), Kir4.1, and aquaporin-4 (AQP-4), that allow astrocytes to maintain synaptic function, extracellular ion homeostasis, and osmoregulation^15–17^. Interestingly, AtAs can occur on a cell-by-cell basis with one astrocyte becoming atypical and losing expression of multiple proteins, while adjacent astrocytes remain normal. AtAs do not express glial fibrillary acidic protein (GFAP), or other intermediate filaments typically upregulated in reactive astrocytes (e.g.: nestin, vimentin), nor do they proliferate^15^. Using the Marmarou weight drop model, previous studies have shown the presence of widespread AtAs in the cortex, with AtAs also noted in the hippocampus. These studies have found that AtAs can arise as early as ten minutes post-injury and persist for at least two months following rmTBI^15, 16^. Here, we investigate whether a mTBI model that includes a single blunt force trauma (BFT) as well as free linear and rotational acceleration forces, induces AtAs and if so, for how long they persist after injury. We employed a closed-head mild TBI (chmTBI) weight drop model in which a blunt force impact to the skull triggered the animal to free fall and land supinely, thus creating a diffuse injury with free linear and rotational motion. We examined an acute timepoint (3-days post-injury) and chronic timepoint (1-month post-injury) after chmTBI. We hypothesized that chmTBI would induce AtAs in areas subjected to the BFT (anterior cingulate cortex [ACC], secondary motor cortex [MOs], retrosplenial cortex [RSC]), as well as in diffuse injury in other brain regions (amygdala, hippocampus, and prefrontal cortex) due to the free linear and rotational nature of the model. We carried out these studies in both male and female mice to determine the effect of sex on AtAs after chmTBI, as previous studies show that sex can influence TBI outcomes^18–21^.

We report that AtAs were abundant 3-days after a single chmTBI in regions subjected to BFT, as well as in the amygdala - an area of diffuse injury. One month after injury, however, the proportion of AtAs between chmTBI and sham injured mice was equivalent in all brain regions. Furthermore, we did not observe significant increases in reactive astrocytes, as traditionally defined by increased GFAP immunoreactivity, after chmTBI in any brain regions. We also found that there was no effect of sex on the abundance or distribution of AtAs after chmTBI. In summary, we report that a single chmTBI, encompassing both a BFT and diffuse injury via free linear and rotational forces, induces AtAs in BFT regions, as well as in the amygdala, without significant induction of reactive astrocytes that can been seen at 3-days, but not 1-month, after injury. Understanding how astrocytes are affected by injury, the presence of reactive and atypical astrocytes, and how this can contribute to downstream neurological and psychiatric dysfunction, will help increase our understanding of TBI pathophysiology and may help to inform specific treatment strategies.

## Materials and Methods

### Animals

All procedures were conducted in accordance with Tufts University Institutional Animal Care and Use Committee (IACUC) regulations. Nine to twelve-week-old C57BL/6 (18–30 grams; bred in house from Jackson Labs or Charles River stocks) and EAAT2-tdTomato mice (24–39 grams)^22^ of both sexes were given food and water *ad libitum*, and kept on a 12-hour light/dark cycle. Animals were randomly assigned to either receive a closed-head mild traumatic brain injury (chmTBI) or sham injury and were sacrificed either 3-days or 1-month (28-31 days) later. Animals were housed in groups of 2–5 mice from the same experimental group.

### chmTBI model

Mice were subjected to a chmTBI using a weight-drop on a closed and unfixed head to allow for free linear and rotational forces^19–21, 23^. The injury apparatus (Figure 1A) stands 185 centimeters (cm) from top to bottom, with a metal box in its center. Three squares of single-ply perforated toilet paper were tightly secured (using clear adhesive tape) across an opening at the top of the metal box to allow for animal placement. The perforation between squares one and two was positioned directly under the site of impact and marked with a circle to aid in the mouse’s head positioning. A hollow metal tube ran from the top of the apparatus to 5 cm above the opening at the top of the metal box and acted as a guide for the 100.5-gram cylindrical steel weight, both of which were approximately 1.3 cm in diameter. A metal wire was placed through a hole in the guide tube to keep the weight in place and was withdrawn to initiate the weight drop. The front side of the metal box was open to allow for injury videography and included a metal guardrail to hold the foam cushion pads in place, which were used to provide the animal with a soft landing. Cuts were made into foam cushions to allow the 100.5-gram cylindrical steel weight to pass through the cavity of the metal box and land in an expanded polystyrene bucket underneath, thereby preventing secondary impact from the weight or subsequent noise that may disrupt or cause further discomfort to the animal.

**Figure 1.**
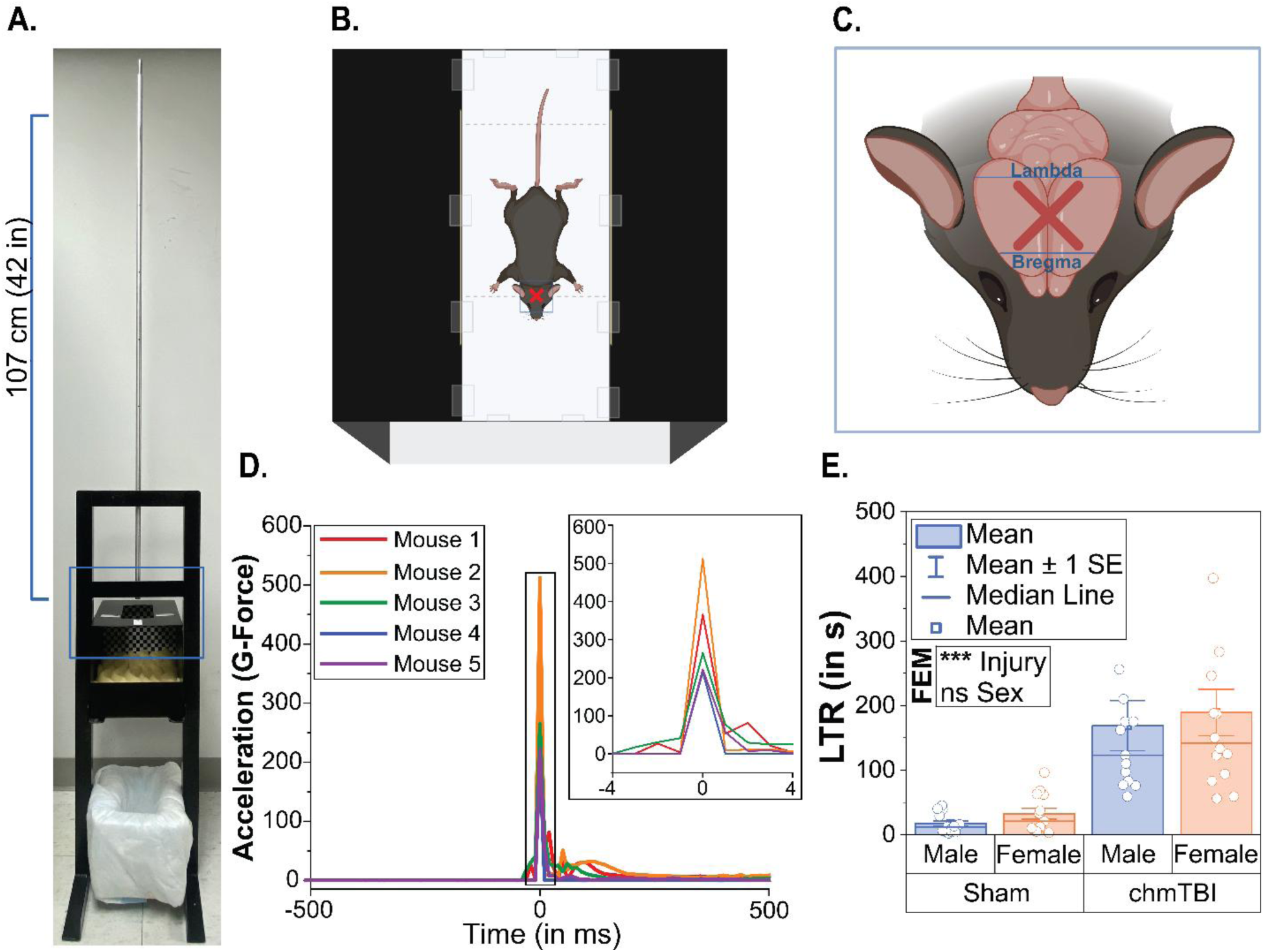
A closed-head mild traumatic brain injury (chmTBI) model with both blunt force and diffuse features. **(A)** The injury apparatus utilizes a 100.5-gram metal rod that is dropped from a height of 107 centimeters. **(B-C)** The site of the rod’s impact is denoted by a red ‘x’. Images made using *Biorender.com*. **(B)** Schematic of mouse positioning prior to weight drop, where the injury is performed on a closed skull. **(C)** The weight is dropped on the midline in between bregma and lambda. **(D)** Representative traces from an accelerometer indicate impacts are consistent with mild traumatic brain injury (mTBI) across trials. Axis-units are represented as acceleration in gravitational force (g-force) and time in milliseconds (y and x-axes, respectively). **(E)** chmTBI male and female mice demonstrate significantly longer latency to right (LTR) than their sham counterparts, thereby indicating an injury-induced effect on consciousness. N = 53 mice, 6–7 animals per condition. Statistical analyses conducted using a fixed effect model (FEM). For all graphs: p<0.05, p<0.01, and p<0.001 indicated by *, **, and *** respectively.

A gravitational force (g-force) range between 200-500 was used to induce mTBI in mice. This represents a force that causes a consistent loss of consciousness after injury without structural damage to the brain. This force accounts for differences in mass scaling and concussion sensitivity in humans^24–27^. The employed g-force range produced no intracranial bleeding, skull fractures, or mortality yet resulted in a significant loss of consciousness as assayed by the latency to righting reflex. Injuries were performed from a drop height of 107 cm. Validation experiments were carried out using an accelerometer (1 kilohertz sampling frequency; 10,000 samples per trial) to ensure consistent impact (Figure 1D). Mice were weighed to ensure they were within the appropriate weight range (see *Animals*), then lightly anesthetized with 3% isoflurane for roughly sixty-seconds before being visually assessed for signs of consciousness. Prior to restoration of the toe pinch reflex, animals were either manually placed on the foam pad below in the supine position (shams), or the weight drop was triggered (chmTBI). Sham animals underwent the exact same procedures as chmTBI animals, minus weight drop impact. Injured animals, referred to as chmTBI, would fall 14 cm before landing in the supine position on the foam pad in the chamber below. Upon waking, mice were returned to their home cage.

### Quantifying latency to right

Latency to right (LTR) was quantified for both sham and chmTBI groups immediately after injury as a measure of injury severity within and between groups. LTR was counted from the time the mouse physically encountered the foam pad until it had turned onto its stomach and demonstrated signs of consciousness. All injuries were video recorded with a stopwatch in the frame to provide timestamps for LTR calculation.

### Immunohistochemistry and Microscopy

Three days or 1-month after chmTBI or sham, mice were deeply anesthetized using isoflurane and assessed for consciousness using the toe-pinch method. Mice were then transcardially perfused using ice-cold solutions of phosphate buffered saline (PBS), followed by 4% paraformaldehyde (PFA). Brains were stored at 4°C and post-fixed overnight in 4% PFA, then switched to 30% sucrose solution for cryopreservation. Tissue was sectioned coronally at 40 μm using a Thermo Fisher Scientific Microm HM 525 cryostat. Following a 1X PBS wash to remove any residual cryopreservation media, tissue was washed in 10% normal goat serum (NGS), 5% bovine serum albumin (BSA), and 0.2% phosphate buffered saline with Triton X-100 (PBS-T) (1:2000 of Sigma-Aldrich Triton^®^ X-100 in 1x PBS) at room temperature on a rocking platform for 1 hour. Sections were then immunolabeled with primary antibodies in a 5% NGS, 1% BSA, and 0.2% PBS-T solution and placed on a rocker at 4 °C overnight. A combination of primary antibodies against GLAST (rabbit; Synaptic Systems 250-113; 1:1,000), GFAP (mouse; Abcam ab7260; 1:500), Kir4.1 (rabbit; Alomone apc-035; 1:200), AQP4 (rabbit; Alomone AQP-004; 1:250), GLT-1 (guinea pig; Millipore Sigma AB1783; 1:1000), and Glutamine Synthetase (rabbit; Millipore Sigma MAB302; 1:100) were used. Slices were washed 3x with PBS, then incubated with chilled secondary antibodies (goat anti-rabbit Alexa 488 [Jackson Immuno; 1:500], goat anti-mouse Alexa 555 [Jackson Immuno; 1:500], or goat anti-guinea pig 647 [Jackson Immuno; 1:500]) at room temperature for two hours on a rocker in a solution of 5% NGS, 1% BSA, and 0.2% PBS-T. Slices were washed twice more with PBS, incubated with 4’,6-diamidino-2-phenylindole (DAPI) for 10 minutes (1:10,000) and then mounted on glass slides. For quantification purposes, brain slices were imaged using an epifluorescence microscope (Keyence BZ-X710) with a 10X dry objective. Excitation and exposure times were consistent across all image collections. For representative images, confocal imaging (Leica SP8) with 20X or 63X oil objectives were used to capture at 0.5–1 µm increments throughout a slice (z-stacks) and subsequently compressed in FIJI^28^ to create a 5–10 µm maximum projection.

### Image Analysis and Quantification

A single experimenter was used for all atypical astrocyte (AtA) and GFAP tissue analysis and was blinded to injury status and sex. To assess the presence of AtAs in C57BL/6 mice, an overlay of GLAST and GFAP was created using FIJI^28^. AtAs were identified as areas lacking GLAST immunoreactivity in distinct clusters with astrocyte-like morphology that did not show upregulation of GFAP immunolabeling. AtAs were manually delineated in FIJI^28^ by drawing a perimeter around each putative AtA and taking the sum within each brain region of interest to estimate the amount of AtAs. The density of AtAs in each area was analyzed as follows: Cortical midline structures subjected to blunt force trauma from the weight’s direct impact (BFT regions) (individually analyzed as Anterior Cingulate Cortex [ACC], Secondary Motor Cortex [MOs], and Retrosplineal Cortex [RSC]), were selected, with each brain region per hemisphere quantified separately for a total of 10-12 BFT regions of interest [ROIs] drawn per animal. The amygdala (inclusive of Lateral Amygdalar nucleus [LA], Basolateral Amygdalar nucleus [BLA], and Medial Amygdalar nucleus [MeA] subregions; images 65-76 from the coronal Allen Mouse Brain Atlas^29^), was selected, with each hemisphere quantified separately for a total of 4-6 amygdalar ROIs per animal. Dorsal hippocampus (inclusive of cornu ammonis (CA) 1, CA2, CA3, and Dentate Gyrus [DG] subregions; images 68-77 from the coronal Allen Mouse Brain Atlas^29^) was selected, with each hemisphere quantified separately for a total of 4-6 hippocampal ROIs per animal. Prefrontal cortex (PFC) (inclusive of Prelimbic [PL] and Infralimbic [ILA] subregions; images 31-41 from the coronal Allen Mouse Brain Atlas^29^), was selected, with each hemisphere quantified separately for a total of 4-6 PFC ROIs per animal. All tissue slices selected for analysis were within a designated range of images from the coronal Allen Mouse Brain Atlas^29^ to ensure comparison of similar brain regions in the neuraxis, and to assist in the manual drawing of ROIs. No distinction was made between cortical or cell layers within each ROI. All data values were expressed as a percentage of AtAs within the total area (each ROI) examined.

The EAAT2-tdTomato transgenic mouse line labels roughly 70-80% of cortical astrocytes^22^. For EAAT2-tdTomato mice, atypical astrocytes were identified as regions that labeled with tdTomato that lacked expression of astrocyte proteins: GLAST and GLT-1, GLAST and GFAP, Kir4.1 and GLT-1, AQP4 and GLT-1, or glutamine synthetase (GS) and GLT-1.

To investigate the presence of reactive astrocytes in C57BL/6 mice, we used CellProfiler 4.2.6 to develop a pipeline that quantified the percent area with GFAP+ astrocytes within an ROI^30^. ROIs were drawn, as above for AtA analysis to create a mask, which the pipeline used as the area to identify putative reactive astrocytes using the adaptive thresholding strategy and Otsu thresholding method. Resulting values were used to calculate the percent area with GFAP+ cells for each animal. The percent area occupied by GFAP+ cells was determined by taking the sum pixel size of primary objects within an ROI mask and dividing by the total pixel size within the ROI mask. Values were then multiplied by 100 to convert into a percentage.

### Statistics

Statistical analysis was conducted using R-Studio 2023.12.1 Build 402. Figures were created using OriginLabs 2025, *BioRender.com*, and Adobe Illustrator. For figures 2-8, large circles represent animal averages, and small circles represent either a single brain region (BFT regions: anterior cingulate cortex [ACC], secondary motor cortex [MOs], or retrosplineal cortex [RSC]), or each hemisphere per slice. Each experimental group had 6-7 mice (3-days: 6 male shams, 6 female shams, 6 male chmTBI, 7 female chmTBI; 1-month: 7 male shams, 7 female shams, 7 male chmTBI, 7 female chmTBI). Normality was tested using the Shapiro-Wilk test.

**Figure 2.**
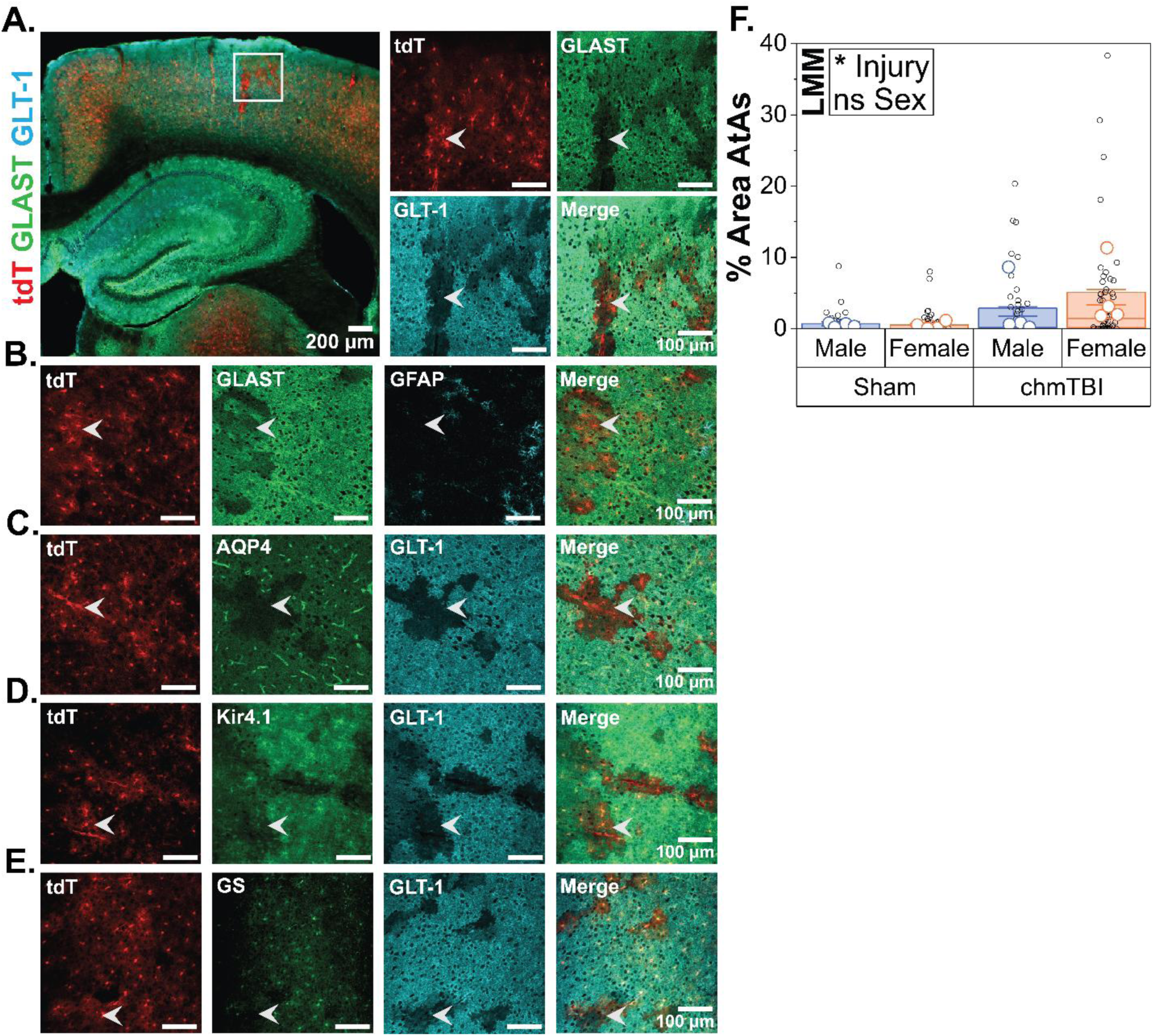
Closed-head mild traumatic brain injury (chmTBI) induces an atypical astrocyte (AtA) phenotype. **(A-E)** EAAT2-tdTomato (tdT) transgenic mouse line is used to label layer II-VI cortical astrocytes (shown in red, leftmost panels). Images taken at 4X and 20X magnification are shown with 200 µm and 100 µm scale bars, respectively. White arrowheads point to example areas with AtAs. Immunolabeling demonstrates colocalized downregulation in regions labeled with tdTomato and were thereby considered AtAs. **(A)** Glutamate/aspartate transporter (GLAST) and Glutamate transporter 1 (GLT-1) **(B)** GLAST and glial fibrillary acidic protein (GFAP) **(C)** Aquaporin-4 (AQP-4) and GLT-1 **(D)** Kir4.1 and GLT-1 **(E)** Glutamine synthetase (GS) and GLT-1 **(F)** Injury has a significant effect on the percent area with AtAs. tdTomato+ areas of astrocyte morphology lacking GLAST expression, and did not have increased GFAP expression, were considered AtAs. N = 16 mice (4 per group, 10-12 regions of interest [ROIs] per animal). Large data points represent animal averages, small data points represent a single brain region (anterior cingulate cortex [ACC], secondary motor cortex [MOs], or retrosplineal cortex [RSC]). Linear mixed modeling (LMM).

Multiple linear regression modeling (MLR) was used to assess differences between sex and injury status for the latency to right behavioral assay. Example code is as follows: C57_3Days_LTR <-lm(Latency_to_Right ∼ Injury_Status + Sex, data = LTR_C573day / summary(C57_3Days_LTR). As seen in Figure 1, we tested whether injury status or sex had a significant difference on latency to right. We employed a multiple linear regression model for this statistical assay to compare differences between the four experimental groups while accounting for the single-measures characteristic of the data.

Linear mixed-effects modeling (LMM) was used for the remaining statistical analyses of timepoint (3-days vs. 1-month), sex (male vs. female), brain region (BFT regions [ACC, MOs, RSC], amygdala, hippocampus, and prefrontal cortex), and injury status (sham vs. chmTBI). We employed LMM to permit use of repeated measures for each mouse without compromising the assumption of independence, as single brain regions (BFT regions: ACC, MOs, RSC), or each hemisphere per slice (amygdala, hippocampus, PFC) were used as the values for our statistical modeling. Dependent variables were the percent area of AtAs or the percent area containing GFAP+ astrocytes. The lme4 package in R-Studio was used to fit the linear mixed model. Example code is as follows: C57_3Days_AtAs <- lmer(Percent_AtAs ∼ Injury_Status + Sex + (1|Mouse_ID), data = C573day_AtAs, na.action = “na.omit”) / summary(C57_3Days_AtAs). Mouse ID was used as the random variable to account for the inherent variability of each mouse.

For all statistical models, alpha values less than 0.05 were considered statistically significant. Statistical significance was denoted using *, **, and *** for p ≤ 0.05, p ≤ 0.01, and p ≤ 0.001, respectively. The relevel function in R-Studio was used to make comparisons within factor levels in lieu of post-hoc testing. Example code is as follows: C573day_AtAs$Sex <- factor(C573day_AtAs$Sex) / C573day_AtAs$Sex <- relevel(C573day_AtAs$Sex, ref = “Male”) / C57AtAs_releveled <- lmer(Percent_AtAs ∼ Injury_Status + Sex + (1|Mouse_ID), data = C573day_AtAs, na.action = “na.omit”) / summary(C57AtAs_releveled). For all statistical models, variables were first run using interaction effects and were thereby fit to a simpler model if presenting with non-significant interaction effects. Any potential outliers were included in analysis, as no statistical testing for outliers was conducted.

## Results

### A closed-head mild traumatic brain injury model with both blunt force and diffuse features

We utilized a chmTBI model which induced a blunt force injury through a weight drop, and a diffuse injury by free linear and rotational motion (Figure 1A-C, see methods for more detail). Mice were lightly anesthetized and placed in a chmTBI apparatus where they were subjected to blunt force trauma (BFT) from a weight drop, followed by free linear and rotational motion which occurred as the mouse fell through an opening and landed in a supine position on padding below. Validation experiments were carried out using an accelerometer to ensure a consistent severity of impact and was delivered between 200 and 500 g-force (Figure 1D). In humans, mTBI can include loss of consciousness (LOC) of up to 30 minutes, with greater lengths of unconsciousness associated with more severe injuries^31^. We measured the animal’s latency to right (LTR) following injury as a behavioral proxy for injury severity and found that mice experiencing a chmTBI had a significantly longer LTR period than mice that underwent sham injury (FEM; t = −3.806, p = 9.67e−4, estimated mean difference = −137.91; Figure 1E). Sex did not affect LTR in chmTBI or sham groups (t = −1.083, p = 0.29, estimated mean difference = −39.24). This suggests that chmTBI had an immediate and significant effect on consciousness, as assayed by LTR.

### chmTBI induces an atypical astrocyte (AtA) phenotype in EAAT2-tdTomato mice

We first carried out chmTBI in EAAT2-tdTomato (tdT) transgenic mice, which contains a bacterial artificial chromosome (BAC) in which tdT is expressed in astrocytes under the control of the EAAT2 promoter. In this mouse line, approximately 70-80% of layer II-VI cortical astrocytes express tdT. Mice underwent the same experimental chmTBI procedure as C57BL/6 and were perfused 3-days later for immunohistochemistry. We examined the immunohistochemical labeling of a panel of astrocyte functional proteins (GLAST, GLT-1, Kir4.1, AQP4, GFAP, and GS) in the cortex of BFT regions.

We found multiple areas that lacked GLAST and GLT-1 immunolabeling but were tdT-positive, thereby demonstrating that despite lacking immunoreactivity of prototypical astrocytic functional proteins, these regions contained astrocytes (Figure 2A). This replicates prior studies^15, 16^ showing that AtAs are abundant in cerebral cortical gray matter within a few days after mTBI or rmTBI. We then asked whether these putative atypical astrocytes were reactive, as classically defined by increased immunolabeling for GFAP. Areas that lacked GLAST, but were tdT-positive, had uniformly low levels of GFAP in the cortex, consistent with AtAs not being reactive (Figure 2B). We further examined whether additional astrocyte functional proteins were lost. Again, we found abundant tdT+ astrocyte regions that lacked Kir4.1, AQP4, GFAP, and GS, similar to previous studies (Figure 2C-E)^15, 16^. Taken together, this supports that AtAs are present in cerebral cortical gray matter 3-days after chmTBI.

Lastly, we quantified the percent area with AtAs in BFT regions (ACC, MOs, RSC) (see Figure 3C for example ROI) and asked if this was affected by injury and sex. We performed immunohistochemistry on sham and chmTBI EAAT2-tdTomato mice 3-days after injury. Regions that were immunonegative for GLAST and GFAP, but were tdT positive (GLAST-/GFAP- /tdTomato+) were considered regions containing AtAs. We found a significant increase in the proportion of AtAs in all BFT regions of chmTBI compared to shams, irrespective of sex (Injury: t = −2.178, p = 0.0481, estimated mean difference = −2.8426; Sex: t = −0.69, p = 0.5020, estimated mean difference = −0.9081; no interaction effect) (Figure 2F). We observed no differences in the percent area containing AtAs between the three BFT brain regions for each experimental group (data not shown).

**Figure 3.**
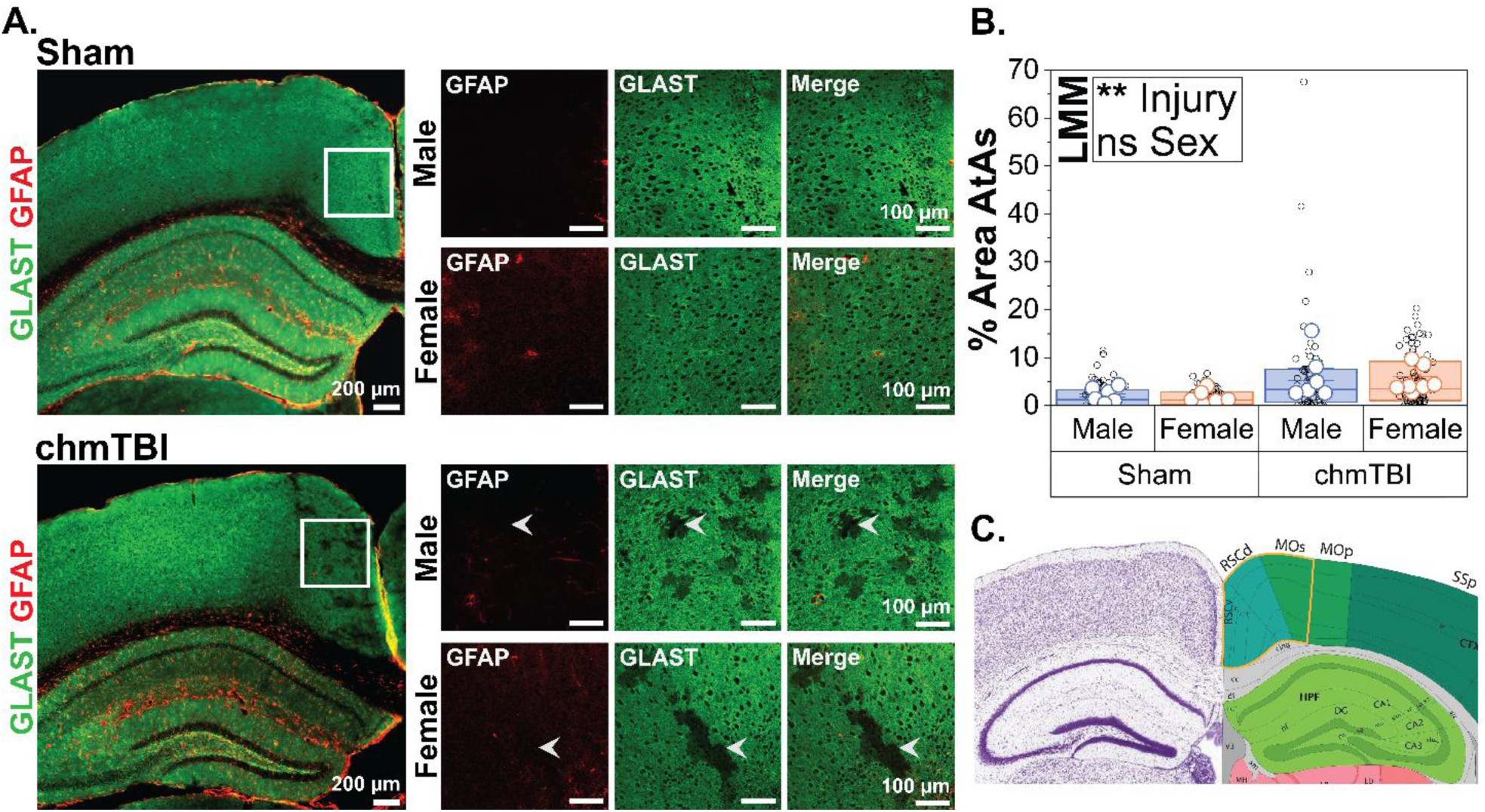
Closed-head mild traumatic brain injury (chmTBI) increases the abundance of atypical astrocytes (AtAs) in blunt force trauma (BFT) regions 3-days post-injury. **(A)** Representative images taken from the retrosplineal cortex (RSC) of sham (top) and chmTBI (bottom) male and female mice, respectively, which were sacrificed 3-days post-injury, immunolabeled with glutamate/aspartate transporter (GLAST, shown in green) and glial fibrillary acidic protein (GFAP, shown in red) and imaged at 20X magnification. Scale bar = 100 µm. **(B)** Injury has a significant effect on AtA presence, but sex does not. The proportion of AtAs was quantified in areas (anterior cingulate cortex [ACC], secondary motor cortex [MOs], and RSC) subjected to BFT from the weight’s direct impact that contained AtAs and was expressed as a percent. No differences were found between brain regions (data not shown). N = 25 mice (6-7 per group, 10-12 regions of interest [ROIs] per animal). Large data points represent animal averages, small data points represent a single brain region (ACC, MOs, or RSC). Linear mixed modeling (LMM). **(C)** Adapted from the Allen Mouse Brain Atlas – Coronal Atlas, at the same position as seen in panel A, *mouse.brain-map.org*, with the region of interest outlined in yellow.

### chmTBI increases the abundance of AtAs in BFT regions 3-days post-injury

After confirming previously published findings that mTBI induces AtAs, we next carried out a more highly powered study using C57BL/6 mice. We quantified the percent area containing AtAs in the BFT regions and again asked if this was affected by injury or sex. We performed immunohistochemistry on sham and chmTBI animals 3-days after injury and considered regions lacking GLAST and GFAP immunolabeling to be containing AtAs. We found a significant increase in the percent area containing AtAs in all BFT regions of chmTBI compared to shams, irrespective of sex (Injury: t = −3.232, p = 0.0038, estimated mean difference = −4.0548; Sex: t = 0.504, p = 0.619, estimated mean difference = 0.6329; no interaction effect) (Figure 3B). We observed no differences in the percent area containing AtAs between the three BFT regions for each experimental group (data not shown).

### chmTBI has no effect on the abundance of AtAs in BFT regions when examined 1-month post-injury

We then asked whether the percent area of AtAs in the BFT regions were affected by injury and sex at the 1-month post-injury time point. We repeated the chmTBI or sham injury protocol on male and female C57BL/6 mice and performed immunohistochemistry 1-month after injury. Again, we considered areas that lacked both GLAST and GFAP to be areas containing AtAs, similar to the 3-days post injury study described as above. We found no difference between injury group or sex in the percent area containing AtAs in the BFT regions (Injury: t = 0.685, p = 0.499853, estimated mean difference = 0.3684; Sex: t = 0.305, p = 0.762718, estimated mean difference = 0.1642; no interaction effect) (Figure 4B).

**Figure 4.**
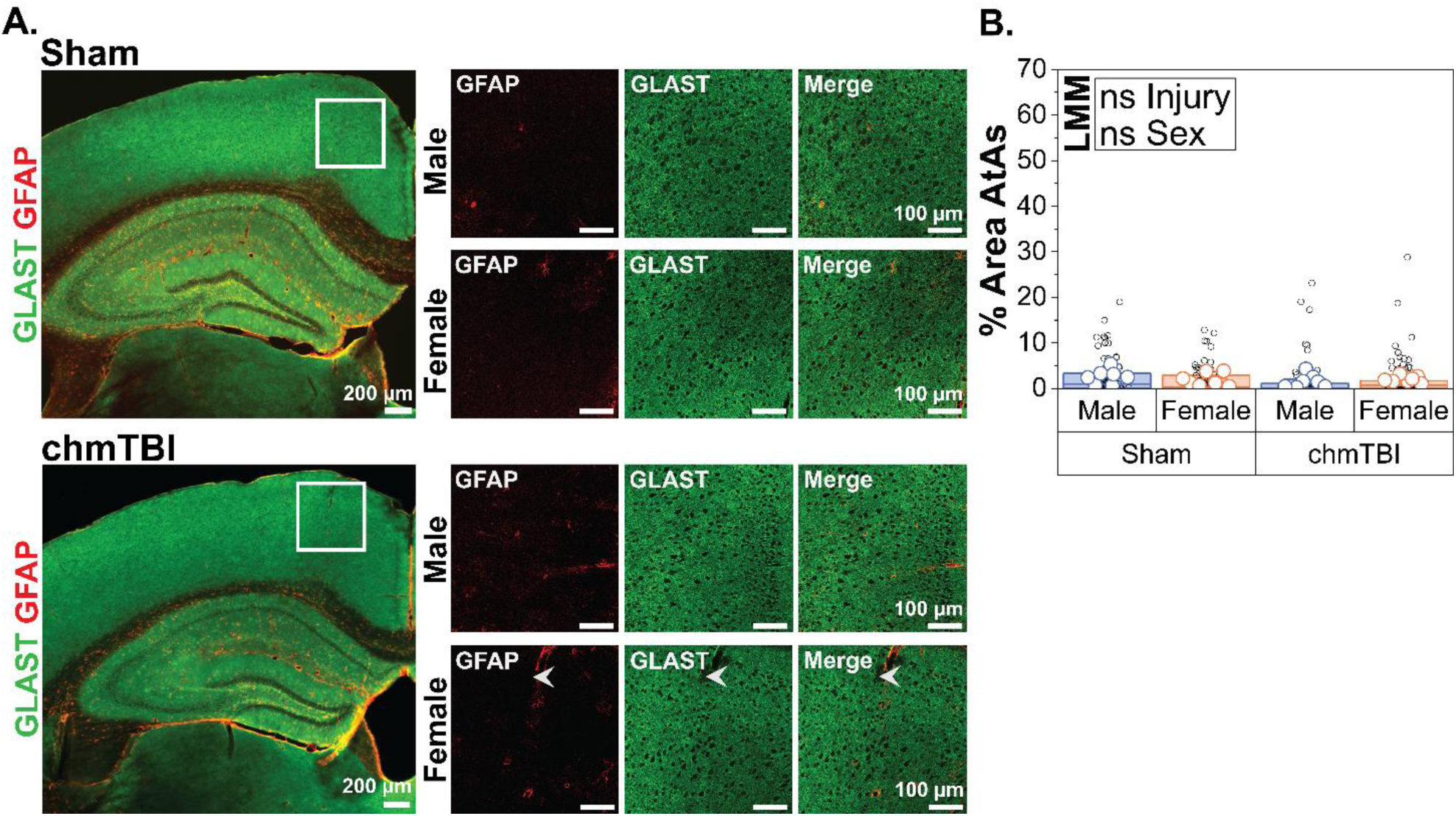
Closed-head mild traumatic brain injury (chmTBI) has no effect on the abundance of atypical astrocytes (AtAs) in blunt force trauma (BFT) regions when examined 1-month post-injury. **(A)** Representative images taken from the retrosplineal cortex (RSC) of sham (top) and chmTBI (bottom) male and female mice, respectively, which were sacrificed 1-month post-injury, immunolabeled with glutamate/aspartate transporter (GLAST, shown in green), and glial fibrillary acidic protein (GFAP, shown in red), and imaged at 20X magnification. Scale bar = 100 µm. **(B)** Neither injury nor sex have an effect on AtA presence. No differences were found between brain regions (data not shown). N = 28 mice (6-7 per group, 10-12 regions of interest per animal). Large data points represent animal averages, small data points represent a single brain region (anterior cingulate cortex [ACC], secondary motor cortex [MOs], or retrosplineal cortex [RSC]). Linear mixed modeling (LMM).

**Figure 5.**
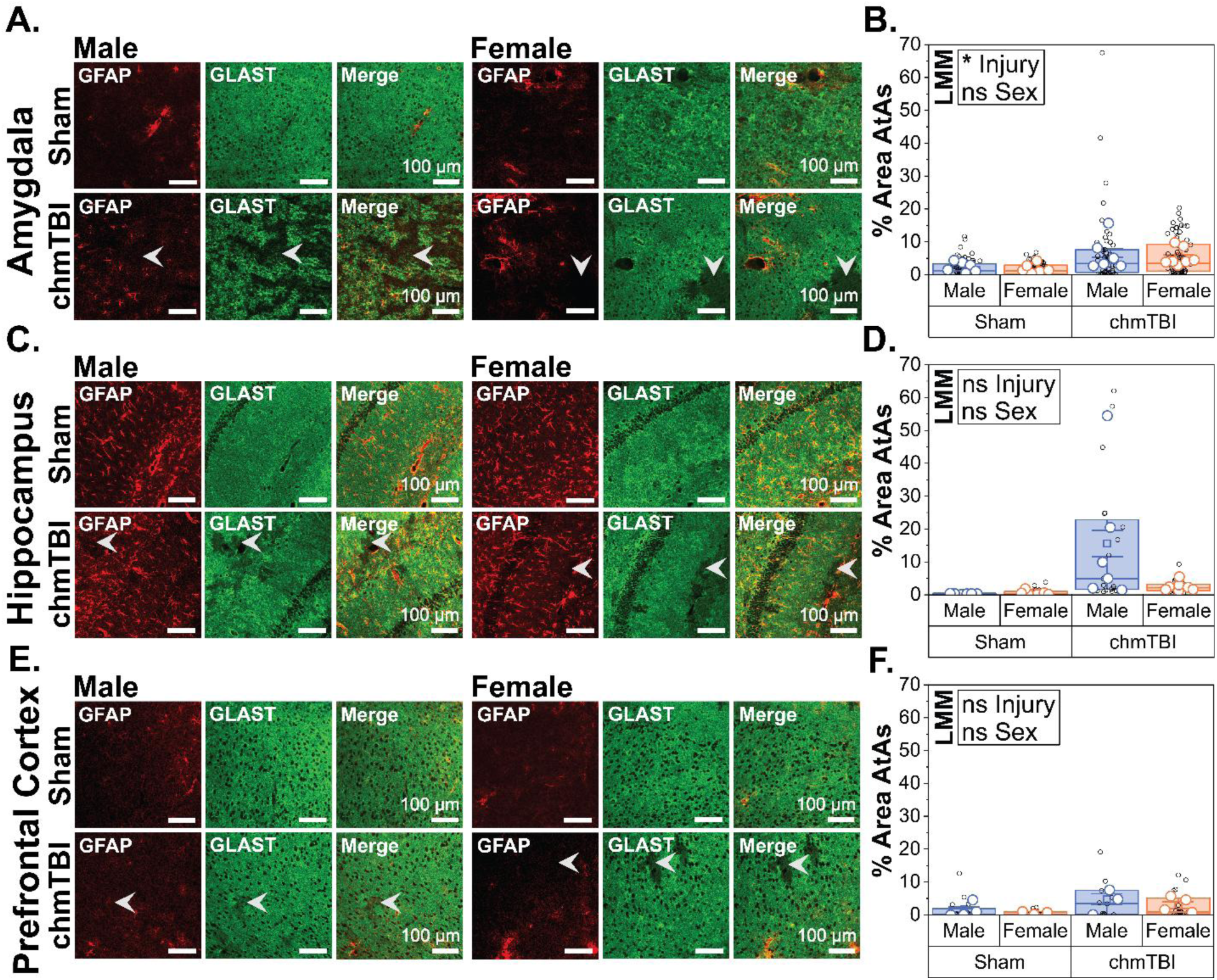
Closed-head mild traumatic brain injury (chmTBI) increases the abundance of atypical astrocytes (AtAs) in the amygdala, but not other diffuse regions, 3-days post-injury. **(A, C, E)** Representative images taken from the basolateral amygdala (BLA), hippocampal cornu ammonis 1 (CA1), and infralimbic prefrontal cortex (ILA), respectively. All representative images taken at 20X magnification, scale bar = 100 µm. **(B)** Injury, but not sex, has a significant effect on the percent area of AtAs in the amygdala (lateral amygdala [LA], BLA, medial amygdala [MeA], subregions pooled). **(D)** Neither injury nor sex have a significant effect on the percent area of AtAs in the hippocampus (cornu ammonis 1-3 [CA1-3], dentate gyrus [DG], subregions pooled), for injury p = 0.0581. **(F)** Neither injury nor sex have a significant effect on the percent area of AtAs in the PFC (ILA, prelimbic cortex [PL], subregions pooled). For amygdala and hippocampus: N = 25 mice (6-7 per group, 4-6 regions of interest [ROIs] per animal), for PFC, N = 13 mice (3-4 per group, 4-6 ROIs per animal). Large data points represent animal averages; small data points represent slice average per hemisphere. Linear mixed modeling (LMM).

### chmTBI increases the abundance of AtAs in the amygdala, but not other diffuse regions, 3-days post-injury

Because the chmTBI model we used includes a diffuse injury component, we investigated brain regions beyond the regions subjected to BFT. Brain sections were co-immunolabeled with GLAST and GFAP, as described above, and regions that were GLAST and GFAP negative were quantified as AtAs. As multiple brain regions can be affected by mTBI, we focused on select brain regions associated with neurological dysfunctions commonly reported following mTBI. We first investigated the presence of AtAs in the amygdala due to its role in emotional regulation and the fact that emotional disruptions, such as mood swings, irritability, anxiety, and depression are commonly reported clinically following mTBI^3, 32^. We found a significant increase in percent area containing AtAs in the amygdala following chmTBI when compared to shams (Injury: t = −2.315, p = 0.0303, estimated mean difference = −7.116) (Figure 4D). Sex did not have a significant effect on the amount AtAs in the amygdala (Sex: t = 1.422, p = 0.169, estimated mean difference = 4.371).

We next investigated the presence of AtAs in the hippocampus, given disruptions associated with learning and memory are commonly reported following mTBI^33, 34^. Overall, neither sex nor injury had a significant effect on the proportion of AtAs within hippocampal regions, despite injury showing a strong trend (Injury: t= −1.997, p = 0.0581 estimated mean difference = −8.284; Sex: t= 1.597, p = 0.1245, estimated mean difference = 6.625). Lastly, we examined the presence of AtAs in the prefrontal cortex (PFC) due to its importance for a variety of complex emotional and cognitive abilities. Following mTBI, emotional and cognitive disruptions typically associated with the PFC are commonly reported clinically, including personality changes, problems with executive functioning, emotional regulation, and impulse control^33, 35, 36^. We found neither injury nor sex had a significant effect on the presence of AtAs in the PFC (Injury: t = −1.959, p = 0.075, estimated mean difference = −2.755; Sex: t = 1.009, p = 0.3342, estimated mean difference = 1.419), although again there was a trend towards an effect of injury.

### chmTBI has no effect on the abundance of AtAs in any diffuse regions 1-month post-injury

We next investigated whether the percent area containing AtAs in diffuse areas (see above) were affected by injury and sex at the 1-month post-injury timepoint. For each brain region, AtAs were quantified using immunohistochemistry, as described above. In the amygdala, we found no significant differences in the percent area containing AtAs between injury or sex groups (Injury: t = 0.237, p = 0.8149, estimated mean difference = 0.40793; Sex: t = −0.02, p = 0.984, estimated mean difference = −0.03501) (Figure 6B). We repeated this investigation in the hippocampus at 1-month post-injury, with no significant differences found in the percent area containing AtAs between injury or sex groups (Injury: t = −0.047, p = 0.96283, estimated mean difference = −0.003148; Sex: t = 0.178, p = 0.86021, estimated mean difference =0.011899) (Figure 6D). Lastly, we investigated the prefrontal cortex at 1-month post-injury and again found no significant differences in the percent area containing AtAs between injury or sex groups (Injury: t = 1.049, p = 0.304, estimated mean difference = 0.2982; Sex: t =0.744, p = 0.464, estimated mean difference = 0.2116) (Figure 6F).

**Figure 6.**
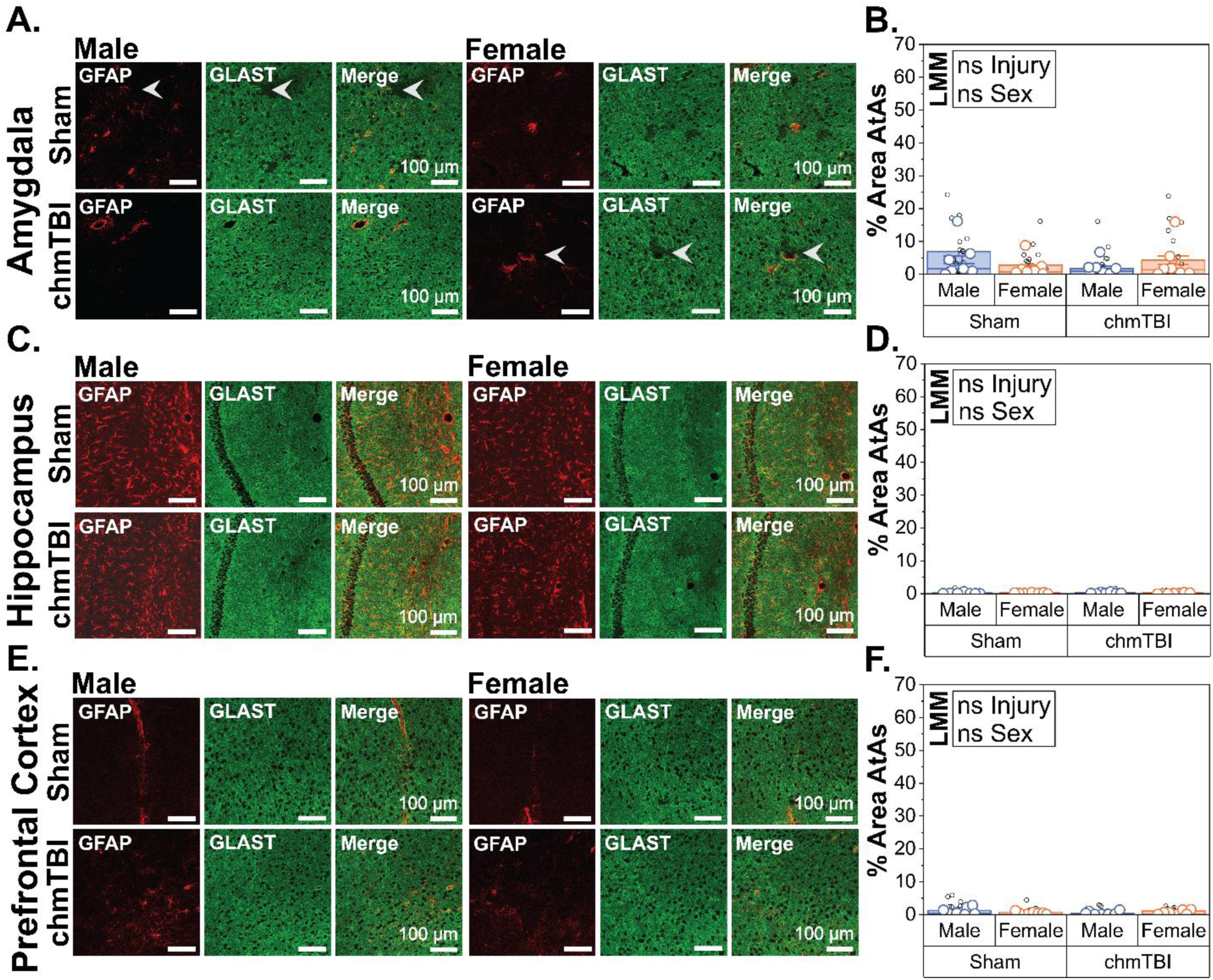
Closed-head mild traumatic brain injury (chmTBI) has no effect on the abundance of atypical astrocytes (AtAs) in any diffuse regions 1-month post-injury. **(A, C, E)** Representative images taken from the basolateral amygdala (BLA), hippocampal cornu ammonis 1 (CA1), and infralimbic prefrontal cortex (ILA), respectively. All representative images taken at 20X magnification, scale bar = 100 µm. **(B)** Neither injury nor sex have an effect on the percent area of AtAs in the amygdala (lateral amygdala [LA], BLA, medial amygdala [MeA], subregions pooled). **(D)** Neither injury nor sex have an effect on the percent area of AtAs in the hippocampus (cornu ammonis 1-3 [CA1-3], dentate gyrus [DG], subregions pooled). **(F)** Neither injury nor sex have an effect on the percent area of AtAs in the prefrontal cortex (ILA, prelimbic cortex [PL], subregions pooled). N = 28 mice (7 per group, 4-6 regions of interest per animal). Large data points represent animal averages; small data points represent slice average per hemisphere. Linear mixed modeling (LMM).

### GFAP immunolabeling is not increased by injury at 3-days or 1-month post-injury

Reactive astrocytes are a common pathophysiological response to TBI and other insults^14^ and in the cortex, reactive astrocytes display increased GFAP immunolabeling. We quantified the percent area of GFAP+ regions in chmTBI and sham animals 3-days and 1-month after injury and examined the effects of injury and sex in all regions mentioned above (BFT regions, amygdala, hippocampus, prefrontal cortex). The percent area containing GFAP+ cells was used as a proxy for the abundance of reactive astrocytes, although this is only one of the many diverse ways astrocytes respond to insult^14^. We found that 3-days post-injury in BFT regions, there were no differences in the percent area occupied by GFAP+ cells, regardless of injury or sex (Figure 7B) (Injury: t = −0.825, p = 0.427, estimated mean difference = −0.4271; Sex: t = 0.693, p = 0.5023, estimated mean difference = 0.3558). In diffuse brain regions 3-days post-injury, there were also no differences observed in the percent area occupied by GFAP+ cells, regardless of injury or sex (Amygdala [Figure 7C] - Injury: t = 0.354, p = 0.7301, estimated mean difference = 0.09142; Sex: t = 0.163, p = 0.8733, estimated mean difference = 0.04172; Hippocampus [Figure 7D] - Injury: t = 0.259, p = 0.8, estimated mean difference = 0.644; Sex: t = −0.62, p = 0.546, estimated mean difference = −1.531; PFC [Figure 7E] - Injury: t = 0.526, p = 0.603, estimated mean difference = 0.37441; Sex: t = −0.017, p = 0.987, estimated mean difference = −0.01179). Taken together, this suggests that a single chmTBI does not induce a significant difference in upregulation of the astrocyte reactivity marker, GFAP, in either sex at 3-days post-injury.

**Figure 7.**
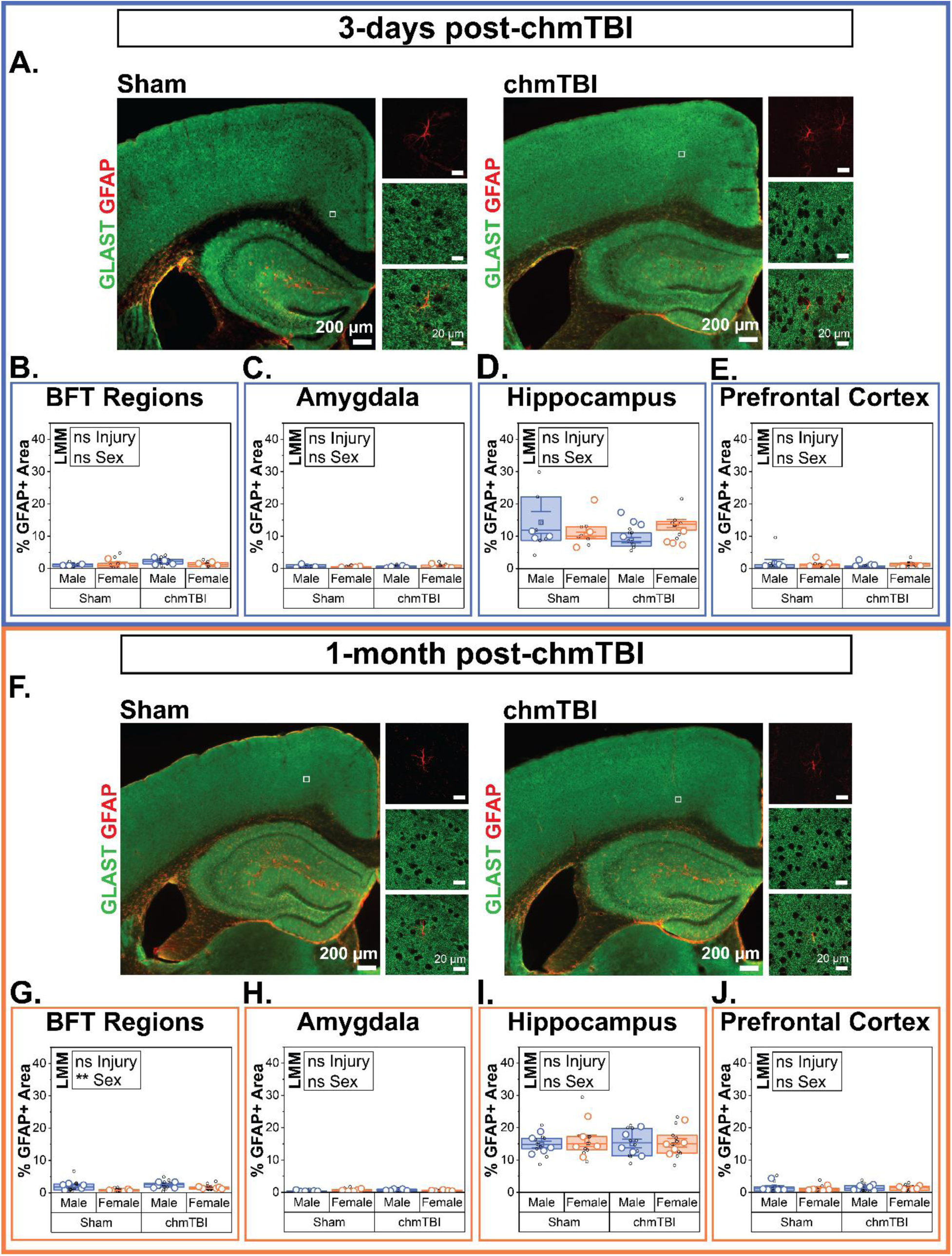
Glial fibrillary acidic protein (GFAP) immunolabeling is not affected by injury at 3-days or 1-month post-injury. **(A, F)** Representative images taken at 4X and 63X magnification in the cortex of sham and closed-head mild traumatic brain injury (chmTBI) animals, which were sacrificed 3-days or 1-month post-injury (left and right boxes, respectively) and immunolabeled with GFAP. For 4X and 63X images, scale bars are 200 µm and 20 µm, respectively. Regions of interest (ROIs) pooled by hemisphere for all graphs. **(B-E)** 3-days post-injury, there are no differences between injury status or sex in the percent area with GFAP immunolabeling for any brain region (blunt force trauma [BFT] regions, amygdala, hippocampus, prefrontal cortex). **(G-J)** 1-month post-injury, there are no injury-induced differences in the percent area with GFAP immunolabeling for any brain regions (BFT regions, amygdala, hippocampus, prefrontal cortex). Sex has an effect on the percent area with GFAP immunolabeling in BFT regions (effect size = 1.0644), but not diffuse regions. N = 34 mice (3-5 per group, 2-3 slices per animal per brain region [BFT, amygdala, hippocampus, PFC). Large data points represent animal averages; small data points represent slice average. Linear mixed modeling (LMM).

We next tested whether there was an effect of injury or sex at 1-month post-injury in the same brain regions (see above). We found that 1-month post-injury in BFT regions, injury had no effect on the percent area occupied by GFAP+ cells, but sex did, with male mice having greater percent GFAP+ area (Injury: t = −2.05, p = 0.05608, estimated mean difference = −0.5885; Sex: t = 3.708, p = 0.00175, estimated mean difference = 1.0644) (Figure 7G). In diffuse brain regions 1-month post-injury, there were no differences observed in the percent area occupied by GFAP+ cells, regardless of injury or sex (Amygdala [Figure 7H] - Injury: t = −0.59, p = 0.563, estimated mean difference = −0.07; Sex: t = −0.062, p = 0.951, estimated mean difference = −0.007366; Hippocampus [Figure 7I] - Injury: t = 0.039, p = 0.969, estimated mean difference = 0.06649; Sex: t = −0.461, p = 0.651, estimated mean difference = −0.78538; PFC [Figure 7J] - Injury: t = −0.164, p = 0.87189, estimated mean difference = −0.06641; Sex: t = 0.63, p = 0.53658, estimated mean difference = 0.25579). Taken together, this suggests that a single chmTBI does not induce sufficient upregulation of the astrocyte reactivity marker, GFAP, at 1-month post-injury.

### AtAs are more abundant 3-days post-injury than 1-month post-injury

Lastly, given significant differences in the proportion of AtAs were observed in select brain regions 3-days post injury between sham and chmTBI animals, yet no differences were observed at 1- month, we analyzed whether time point influenced the proportion of AtAs. All values obtained from the aforementioned analyses (Figures 3-6) were utilized, with brain region and sex pooled. A significant increase was found in the percent area containing AtAs at 3-days post-injury compared to 1-month post-injury (t = 2.762, p = 0.0104, estimated mean difference = 5.132) (Figure 8B). This supports the idea that following a single chmTBI, AtAs are transiently present after injury.

**Figure 8.**
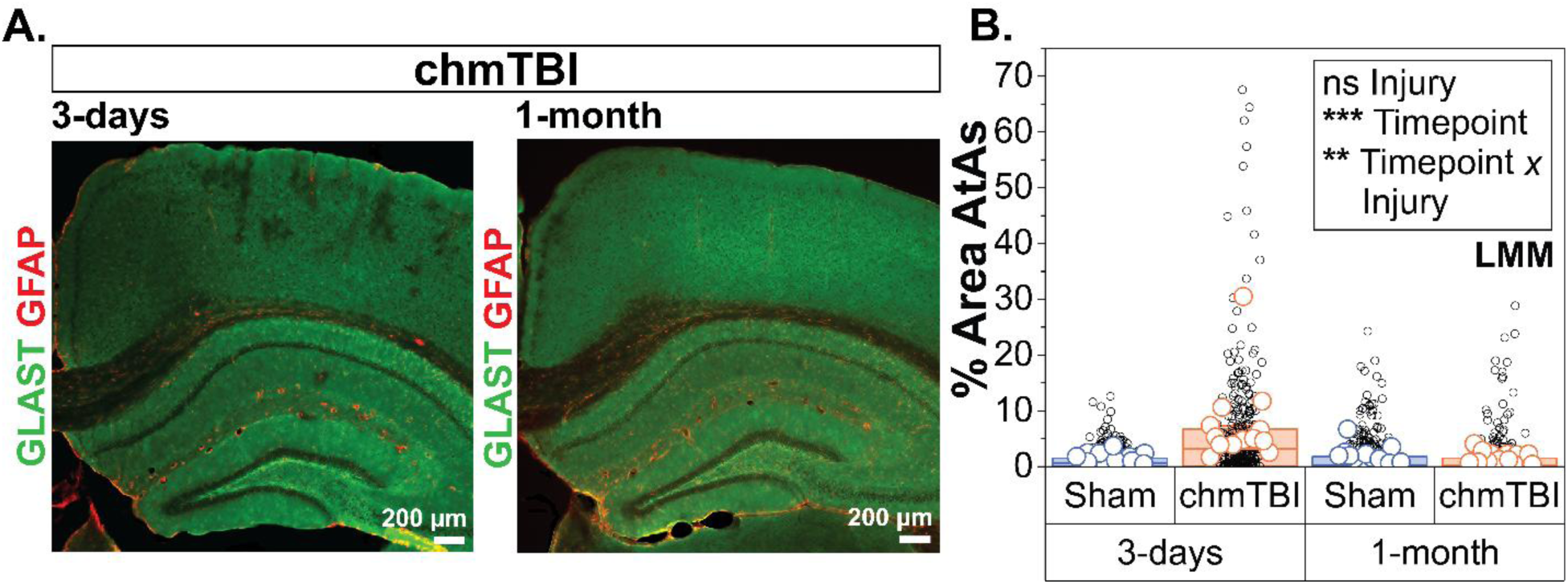
Atypical astrocytes (AtAs) are more abundant 3-days post-injury than 1-month post-injury. **(A)** Representative images taken at 4X in the cortex of closed-head mild traumatic brain injury (chmTBI) animals at 3- days and 1-month post-injury (left and right boxes, respectively), and immunolabeled with glutamate/aspartate transporter (GLAST, shown in green) and glial fibrillary acidic protein (GFAP, shown in red). Representative images taken at 4X magnification, scale bar = 200 µm. **(B)** While there is no significant effect of injury, there is a significant timepoint effect and a significant interaction effect between timepoint and injury (effect sizes 5.225 and 5.266, respectively). Sex and brain region (blunt force trauma [BFT] regions, amygdala, hippocampus, and prefrontal cortex) are pooled. N = 53 mice (12-14 animals per group). Large data points represent animal averages, small data points either represent a single brain region (for BFT regions) or the slice average per hemisphere (for amygdala, hippocampus, PFC) with 18-28 data points per animal. Linear mixed modeling (LMM).

## DISCUSSION

Here we tested whether a chmTBI with BFT and free linear and rotational motion induces AtAs at an acute (3-days) and chronic (1-month) timepoint following injury. We also examined if there was an effect of sex, and whether this injury model induced AtAs in BFT regions (ACC, MOs, RSC) and diffuse regions (amygdala, hippocampus, PFC). We found that 3-days after injury in BFT regions, there was a significantly greater percentage of AtAs in chmTBI animals compared to shams, without induction of a GFAP-positive reactive phenotype. Three-days post-injury in diffuse brain regions, chmTBI induced a significantly greater percentage of AtAs in the amygdalae of injured animals compared to shams, but no significant differences were observed in the hippocampus or PFC. No sex differences were observed in the abundance of either AtAs or GFAP-reactive astrocytes for any brain regions. We found that 1-month after injury in BFT and diffuse brain regions, there were no differences in the percent AtAs between sham and chmTBI animals, or between sexes. Similarly, no differences were observed 1-month post-injury in the percent area with GFAP-positive astrocytes for BFT or diffuse regions, though a small, yet statistically significant, difference was observed between sexes in BFT regions, with males demonstrating a greater percent area with GFAP-positive astrocytes.

Existing AtA studies have focused on rmTBI in a diffuse model which employed limited linear- rotational forces and reported that AtAs persisted as long as two-months post-injury^15^. Here, we report a chmTBI model results in AtAs 3-days post-injury in the cerebral cortex using a model that incorporates both BFT and free linear and rotational motion. As only one study to date has made note of AtAs in the hippocampus following mTBI^15^, we expanded upon this by formally quantifying hippocampal AtAs, in addition to exploring the amygdala and prefrontal cortex regions. While previous studies observed hippocampal AtAs in a modified Marmarou model of repeated injury^15^, we do not see a significant increase in the hippocampus at 3-days post-injury or 1-month post-injury, although there is a strong trend toward AtAs in injured animals 3-days post-injury. This could be due to the model itself, but more likely, due to the repeated versus single injury approaches. Since the astrocyte proteins that are lost in AtAs are important to their function, we suspect AtAs may be associated with disrupted astrocyte glutamate uptake, K^+^ buffering, and compromised BBB. Disrupted BBB has been seen in association with AtAs in a previous study^16^. Given that we and others report AtAs in the cortex and/or amygdala, it is possible they contribute to behavioral disruptions associated with these regions. In future work, we hope to examine astrocytic, neuronal, and circuit-level dysfunction in regions where chmTBI induces AtAs.

In line with the existing AtA literature, we did not observe significant sex differences in AtAs after mTBI, supporting that AtA induction is not sex-specific in this context. However, given AtA research is still emerging, and sex-specific microglial and neuronal changes have been demonstrated after injury^18^, future sex-balanced studies would continue to be of value to the field. With the exception of one study^19^, research using this rotational mTBI paradigm has used male mice^20, 21^, and the former did not report any sex-related differences in allodynia following injury^19^. Further mixed-sex studies will help determine whether there are any sex-specific changes that occur after mTBI in this model. The existing AtA literature has demonstrated astrocyte reactivity for up to 14-days following a modified Marmarou model of repeated injury^15, 16^, in which sex differences were noted at various time points. While differences in reactive astrocytes were not observed in our study following a single chmTBI, we suspect that a repeated injury paradigm would result in astrocyte reactivity. In addition, ‘reactive’ astrocytes are an extremely broad term, meaning AtAs could represent a GFAP-negative form of reactive astrocytes^14^. Past studies have suggested astrocyte reactivity is a graded response to injury severity, which if true, would suggest that our employed injury paradigm is milder in nature^14^. However, given the chmTBI model used here has yet to be studied in a repeated context, this will be a necessary step to elucidate the relationships between these two injury models. Whether injury severity coincides with both increased atypical and reactive astrocytes, and to what extent, would be an interesting topic for future research. In addition, understanding the injury-related disruptions that lead to AtAs, versus reactive astrocytes, would greatly help to understand how injury affects astrocyte gene and protein regulation, astrocyte function, and inflammatory signaling. Investigating the temporal and spatial dynamics between atypical and reactive astrocytes will also be a valuable step toward understanding the interplay between these two astrocyte subtypes that arise following TBI.

One limitation of this study is that the estrous cycle was not tracked. GFAP levels in the amygdala^37, 38^, hippocampus^39, 40^, and neocortex^41^ fluctuate with the estrous cycle under physiological conditions, but no studies have analyzed its changes in response to mTBI at different stages of the estrous cycle. Furthermore, estrogens, which fluctuate during the different estrous cycle stages, can regulate the expression of BBB proteins like Claudin-5^42, 43^. Since an AtA response is thought to be triggered by BBB disruption and astrocytes regulate BBB function^16^, it would be important to check whether the stage of the estrous cycle could influence the percentage of astrocytes that engage in this response. Finally, previous studies in patients suggest that the stage of the menstrual cycle can predict the severity of post-mTBI symptoms^44^. Yet, whether astrocytes play a role in these differences across the menstrual cycle remains unstudied. For all these reasons, documentation or control of the estrous cycle at the time of injury for future studies could be a valuable parameter to explore.

### Conclusion

Our findings provide insight into astrocytic changes following TBI, which may aid in the field’s understanding of how secondary injury unfolds in relation to neurotransmitter homeostasis and other astrocyte functions. Here we employ a closed-head weight-drop model of mild TBI in a sex-balanced cohort to investigate atypical and reactive astrocytes in BFT and diffuse brain regions at an acute and chronic timepoint following mTBI. Our results indicate that our chmTBI model induces AtAs in regions subjected to BFT, and in some diffuse regions, when examined 3-days post-injury; but that the injury-induced increases in AtAs do not persist at the 1-month time point. This suggests AtAs induced by this injury model may be transient, and that by 1-month post-injury AtAs may regain the expression of key homeostatic proteins.

### Transparency, rigor, and reproducibility

Mice (9–12 weeks old) were randomly assigned to sham or chmTBI groups, with 4–7 mice per sex, injury status, genotype, and timepoint groups. All AtA and GFAP tissue analyses were conducted by the same experimenter, who was blinded to injury status and sex; timepoint could not be blinded due to extensive temporal separation between tissue collection and analysis of the 3-days and 1-month cohorts. Analysis of AtAs was performed manually in FIJI^28^ using lab-established protocols. For all analyses, ROIs were drawn manually in consultation with the coronal Allen mouse brain atlas^29^. All statistics were calculated using R, and an alpha level of 0.05 was used for significance testing. LMM was chosen to accommodate the repeated-measures data structure without violating assumptions of independence. Formal power analysis was not conducted; however, sample sizes were designed to maximize scientific and statistical rigor in alignment with field standards. While sample sizes were largely consistent, a smaller sample size was used for the automated GFAP analysis due to vascular artifacts caused by cross-reactivity of the mouse-derived GFAP antibody in animals with suboptimal perfusions. Additionally, a small but notable reduction in PFC sample size for animals sacrificed 3-days post-injury occurred due to insufficient collection of anterior brain tissue during cryosectioning. See *Materials and methods* for further details regarding statistical and experimental analysis.

## Acknowledgements

We thank past and present members of the Dulla lab, in addition to Drs. Kenneth Amaya, Camila Demaestri, Kaitlyn Dorst, Donald Katz, Raymond Knight, Sonja Krstic, and Grant Weiss for their mentorship, guidance, feedback and support.

## Author’s Contributions

**Jesse B. Blackman** (Data curation, Formal analysis, Investigation, Project administration, Software, Supervision, Visualization, Writing – original draft), **Rebecca Krauss** (Data curation, Formal analysis), **Sadi Quiñones** (Software, Supervision), **Panorea Tirja** (Validation), **Mary E. Sommer** (Validation), **Carmen Muñoz-Ballester** (Writing – review & editing), **Farzad Noubary** (Resources, Software), **Moritz Armbruster** (Supervision, Writing – review & editing), **Stefanie Robel** (Writing – review & editing), **Trent Anderson** (Methodology, Resources, Software, Writing – review & editing), **Chris. G. Dulla** (Funding Acquisition, Methodology, Supervision, Writing – review & editing).

## Funding Information

This work was supported by the National Institute on Aging (R01AG085666 and R21AG072905), in addition to the Ellison foundation.

## Author Disclosure Statement

No competing interests, personal financial interests, funding, employment, or any other competing interests exist.

## Notes

### Competing Interest Statement

The authors have declared no competing interest.

